# SVDquintets: a New Tool for Species Tree Inference

**DOI:** 10.1101/2022.06.01.494348

**Authors:** A. Richards

## Abstract

Species tree inference is complicated by the fact that different segments have the genome can have evolutionary histories that differ from each other and from the evolutionary history of the species as a whole. One source of this mismatch is incomplete lineage sorting (ILS), which is commonly modeled by the multispecies coalescent process. Here we derive site pattern probabilities under the multispecies coalescent model, the Jukes-Cantor substitution model, and a relaxed molecular clock for five species at a time. As a result, we can demonstrate that the rank results that form the theoretical basis for SVDQuartets also appear to hold for sets of five species. Based on this, we have developed a new species tree inference algorithm: SVDquintets. Comparison with SVDQuartets shows improved species tree inference under a variety of simulated data settings.

## 2 Introduction

We describe the evolutionary histories of the species via a species tree:

### Definition 1.

A *species tree* is an acyclic graph *S* = (*V* (*S*), *E*(*S*), ***τ*** _*S*_) where *V* (*S*) is the vertex set of *S, E*(*S*) is the edge set of *S*, and ***τ*** _*S*_ is a vector of node times.

The external nodes on the species tree represent extant species while the internal nodes represent speciation events. Internal branches represent a continuum of ancestral species. If we know the common ancestor of all the species under consideration, then the tree is *rooted*, and the tree becomes a directed graph from the root outward. We will assume for our probability calculations that the root of the species tree is known. We also make the simplifying assumption that each speciation event results in two daughter species, so all our trees are binary trees.

Species tree inference is complicated by the fact that the evolutionary history of a species and individual sections of the genome may not match. Causes for this divergence include incomplete lineage sorting (ILS; commonly modeled by the coalescent process [8]), gene duplication and loss (GDL) and horizontal gene transfer (HGT). The history of individual loci on the genome is represented by a gene tree. We emphasize that in the sequel, in keeping with common phylogenetic usage, “gene” refers to a recombination-free sequence of nucleotides, and this may not correspond with the biological meaning of a gene that refers to a segment of DNA that codes for a polypeptide.

### Definition 2.

A *gene tree* is an acyclic graph *G* = (*V* (*G*), *E*(*G*), ***t***_*G*_) where *V* (*G*) is the vertex set of *G, E*(*G*) is the edge set of *G*, and ***t***_*G*_ is a vector of node times. These node times are subject to the constraint that the coalescent events between a set of species must occur prior to the speciation time of the species in question.

We assume that we have one individual sampled from each species and no missing data, so both trees have a common leaf set *L*. When necessary to distinguish the leaves of *G* from the leaves of *S*, we use capital letters *A* through *E* for the leaves of the gene tree and lower case *a* through *e* for the leaves of the species tree. Internal nodes on the gene tree represent coalescent events, which identify the most recent common ancestor of two gene lineages.

A number of different approaches have been taken with regard to species tree inference in the presence of ILS. The first is essentially to ignore the problem: perform gene tree inference on concatenated data using methods such as RAxML [24], FastTree [16], or IQ-TREE 2 [12]; treating all sites as if they share a single, common evolutionary history. This can be fast and accurate for estimating *S*. But, there are some concerns: concatenation has been shown to be statistically inconsistent for some values of (*S*, ***τ***) [3, 21], and speciation time estimates are biased since the coalescent event must naturally occur before the speciation time. Another approach is the use of summary statistics that first estimate the gene trees independently for each gene, and then use the gene tree estimates as inputs for species tree estimates. Examples of this approach include STEM [9], ASTRAL [13, 14, 28], and MP-EST [11]. These methods can be computationally efficient, however they depend on the accuracy of the gene tree estimates that are used as inputs as well as proper delineation of recombination-free segments of the genome [6, 23]. A third approach uses the full data to coestimate the species tree and each of the gene trees, generally using Markov chain Monte Carlo (MCMC) methods. Examples of this approach include BEST [10], *BEAST [4, 15], and BPP [27]. These methods can be quite accurate but are very computationally intensive when the number of loci and/or leaf set is large. Assessment of convergence is also a challenge, especially due to the multi-modal nature of the likelihood in the tree space [22].

A fourth approach, and the one we take in this paper, is to treat the gene trees as a nuisance parameter that can be integrated over. This assumes that each nucleotide has its own history, which is a random draw from the distribution of all possible gene trees given the species tree. Previous examples of this approach include SVDQuartets [1] and Lily-Q [18]. SVDQuartets takes site pattern frequencies and forms a flattening matrix (described in greater detail later) for each possible 4-taxon unrooted species tree. The species tree inferred is based upon the estimated rank of the associated flattening matrix. Lily-Q instead takes the site pattern probabilities and then assumes a distribution on speciation times and the species tree topologies to find a posterior probability of the species tree topology given the data. This was shown to be fast and highly accurate, but the site pattern probabilities were calculated assuming a strict molecular clock. Inference using Lily-Q was very sensitive to this assumption, prompting the need to generalize the site pattern probabilities to cases for which the clock does not hold [19]. This paper further generalizes this work for four taxa to site pattern probabilities for five taxa.

## 3 Model

Our model is as follows. We assume that there is no missing data and that each site is properly aligned across species and has an independent evolutionary history from every other site given the species tree. In other words, each site has its own gene tree which is a random draw from the distribution of gene trees *P* (*G* = *g*|*S*). We also assume neutral selection, which implies that the substitution process and coalescent process are independent [26]. The species tree topology is given by *S* and any given gene tree topology is given by *G*. ***τ*** refers to a set of speciation times and ***t*** a set of coalescent times. *θ* = 2*N*_*e*_*μ* refers to the population parameter, where *N*_*e*_ is the effective population size and *μ* is the mutation rate per generation. Time is measured in coalescent units, or 2*N*_*e*_ generations, so one unit of time is the expected time until a coalescent event between any two lineages. We assume a constant *θ* throughout the tree, because previous work has found that inference is not strongly dependent on *θ* over a biologically plausible range of values [18] and allowing both *θ* and the mutation rates to vary over time would result in a non-identifiable model.

Relative mutation rates are given by *γ*_*ij*_. The index *i* refers to the branch of the tree, labeled by the species descending from the branch. So, *i* = *a* refers to the pendant branch leading to species *a, i* = *bc* would refer to the common branch above the speciation event between *b* and *c*, and so on. The branch above the root is given by *i* = *r*. The index *j* describes the ordered relative rates from the present day back toward the root. All rates are measured relative to the final rate above the root 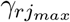. We assume the rates are piecewise constant with the set of breakpoints for the rates given by *ψ*_*ij*_ with *γ*_*ij*_ being the rate between 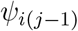 and *ψ*_*ij*_.

Our goal is to estimate the vector of site pattern probabilities given the species tree, speciation times, the mutation rates and the population parameter. For example, let *δ*_*T CCGT*_ = *P* (*i*_*a*_ = *T, i*_*b*_ = *C, i*_*c*_ = *C, i*_*d*_ = *G, i*_*e*_ = *T*) where *i*_*x*_ is the state for species *x*. Under the Jukes-Cantor substitution model, *δ* _*T T T T T*_ = *δ*_*CCCCC*_ for example, so the vector of site pattern probabilities can be mapped down to a 51 × 1 vector ***δ*** = *δ*_*XXXXX*_, *δ*_*XXXXY*_, … *δ*_*Y ZW XX*_. The 51 possible site patterns are listed in table 1.

**Table 1:**
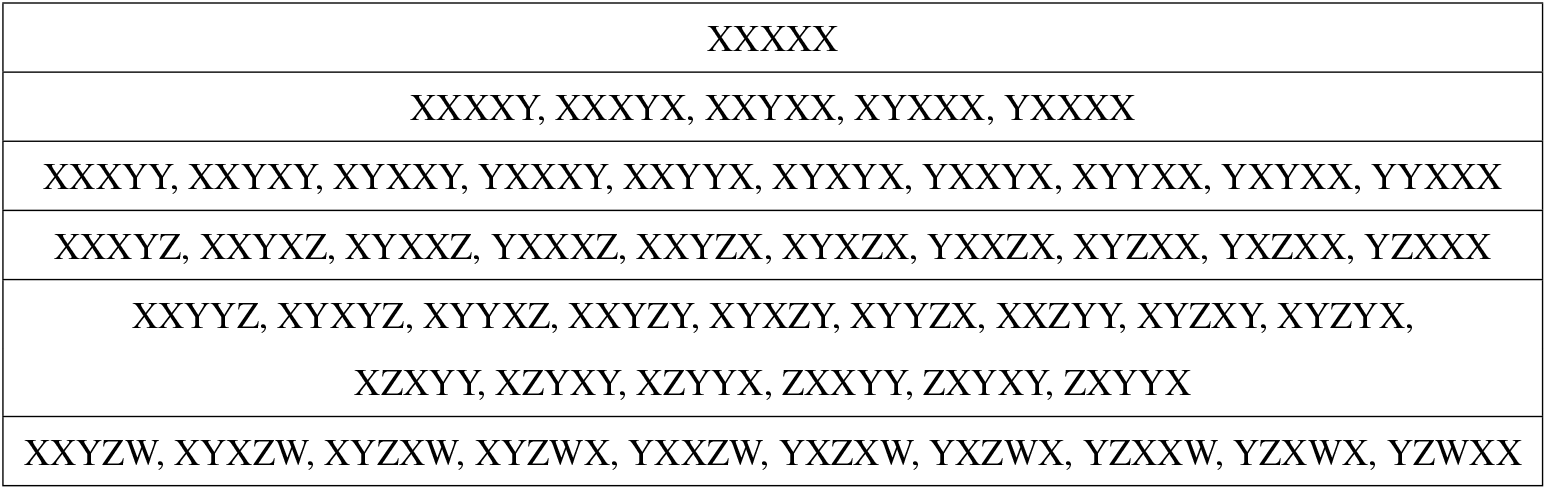
Unique 5-taxon site patterns under the Jukes-Cantor mutation model

The model is summarized by figure 2. Here we show an example species tree with the topology (*a*, ((*b, c*), (*d, e*))). There are two different rates on the common branch above *d* and *e* and three different rates on the common branch above *b* and *c*. All other branches have a single rate and the rate above the root is assumed to be unity (when time is measured in mutation units). One example gene tree is shown embedded within this species tree. In this case the gene tree has a different topology than the species tree: ((*A, B*), (*C*, (*D, E*))). Note that in this case, the coalescent event at time *t*_1_ occurred on the first branch above the (*d, e*) speciation event, but it could have occurred higher up in the tree; likewise the coalescent event at time *t*_2_ could also have occurred both at its actual location or above the root.

**Figure 1:**
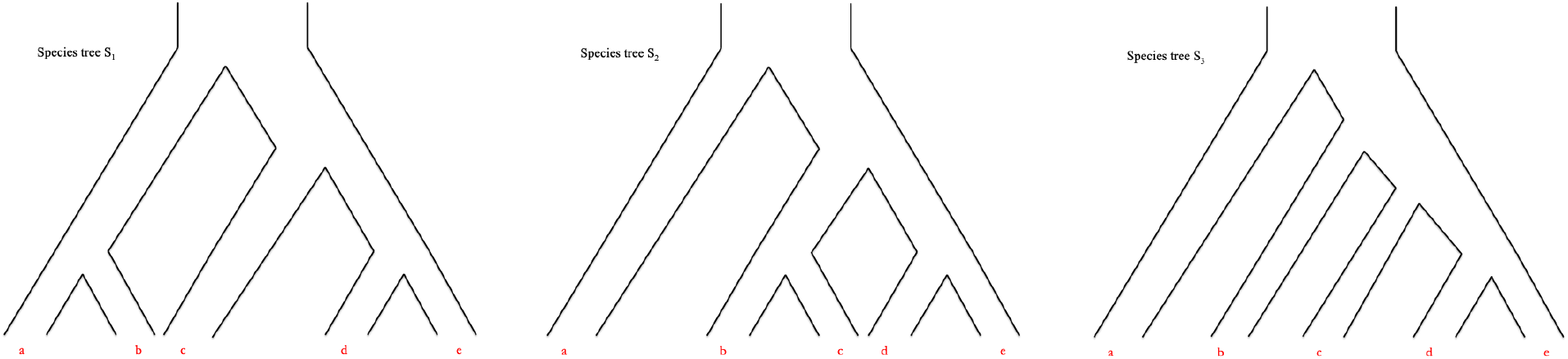
The tree model species tree topologies. The tree model gene tree topologies have the same basic shape as these.

**Figure 2:**
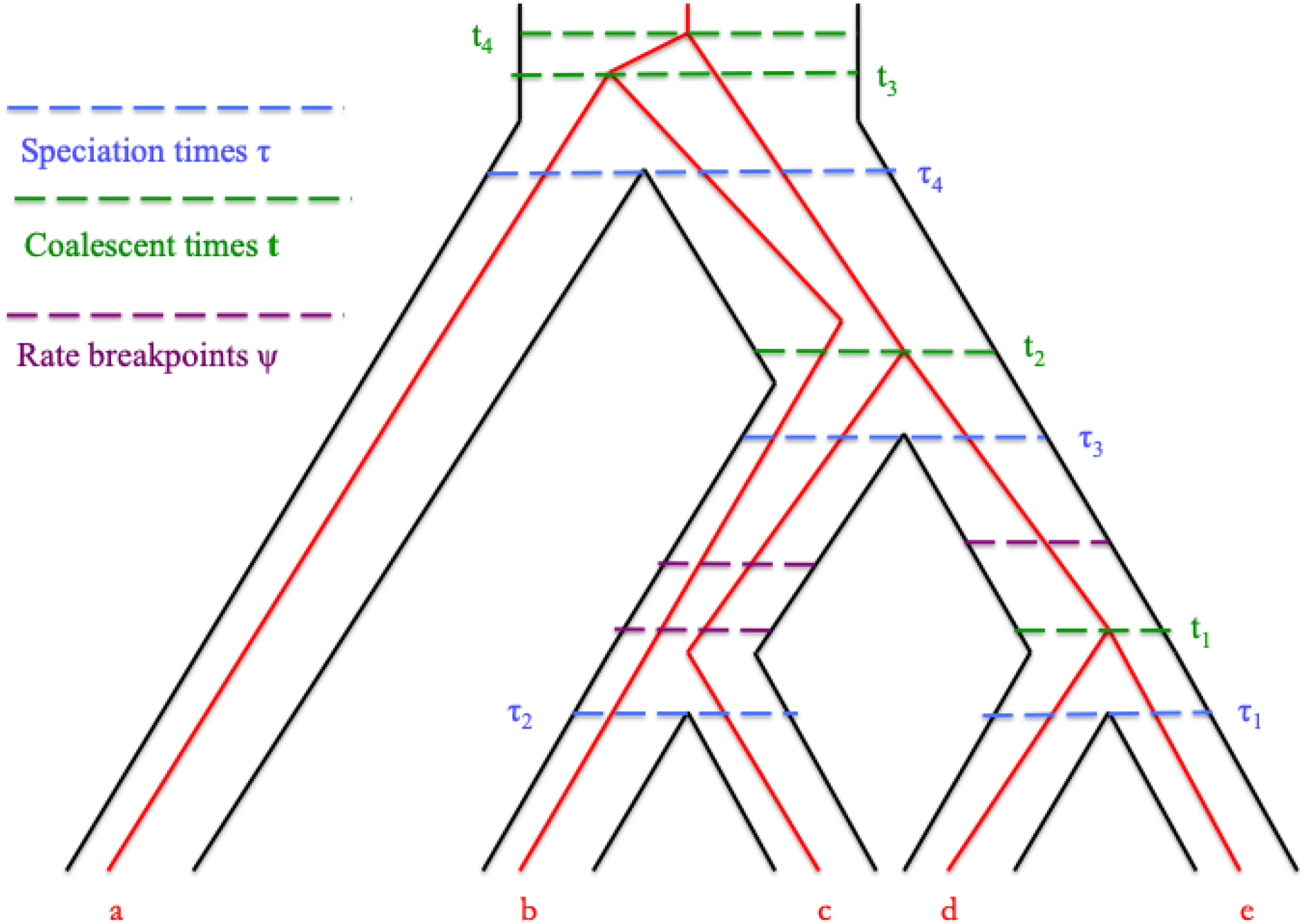
An example gene tree embedded in a species tree. Here the species tree has topology *S*_2_ = (*a*, ((*b, c*), (*d, e*)) while the gene tree has topology *G*_1_ = ((*A, B*), (*C*, (*D, E*))). There are two rates on the branch DE and three on the branch BC, while all other branches evolve at a single rate.

Since we view the gene trees as a nuisance parameter, to compute the site pattern probabilities we integrate over all gene trees consistent with the species tree, G_*S*_:

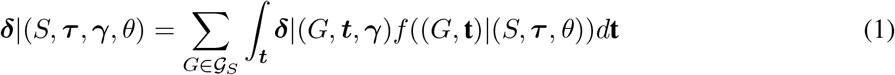

We will discuss the derivation of both of these terms beginning with the gene tree density given the species tree.

### 3.1 Preliminaries

So far, we have only mentioned three species tree topologies of concern, *S*_1_ = ((*a, b*), (*c*, (*d, e*))), *S*_2_ = (*a*, ((*b, c*), (*d, e*))), and *S*_3_ = (*a*, (*b*, (*c*, (*d, e*)))). The reason is that the difference between, for example, *S*_1_ and ((*c, e*), (*d*, (*a, b*))) is only one of labeling. Although there are in total 105 different *labeled* 5-taxon binary tree topologies, there are only these three *unlabeled* topologies (see the lengthy discussion in [7]). All site pattern probabilities given some other topology than *S*_1_, *S*_2_, or *S*_3_ can be derived by permuting the states at the tips of one of these three trees. For example, *δ*_*XXXXY*_ |((*a, b*), (*c*, (*d, e*))) = *δ*_*Y XXXX*_ |((*e, b*), (*c*, (*d, a*))). Similarly, even though there are 105 possible gene tree topologies, we we need only concern ourselves with *G*_1_ = ((*A, B*), (*C*, (*D, E*))), *G*_2_ = (*A*, ((*B, C*), (*D, E*))), and *G*_3_ = (*A*, (*B*, (*C*, (*D, E*)))) after accounting for permutations. For *G*_3_ the coalescent events must occur in a particular order, but for *G*_2_ either *t*_1_ *< t*_2_ or *t*_1_ *> t*_2_ could occur, and with *G*_1_ we can have *t*_3_ *< t*_1_, *t*_1_ *< t*_3_ *< t*_2_, or *t*_2_ *< t*_3_, resulting in 180 different combinations of topology and coalescent time orders.

Not all gene tree phylogenies (*G*, ***t***) are possible given (*S*, ***τ***). For example, given *S*_3_ and *G*_1_, *t*_1_ can occur any time before *τ*_1_, but *t*_3_ must occur above the root. Table 2 summarizes the different cases for what combinations of coalescent events are possible on different branches of the tree for *S*_3_. Similar tables can be constructed for *S*_1_ and *S*_2_.

**Table 2:** Possible coalescent times ***c*** embedded within the species tree *S*_3_. For *G*_1_, there is a direct correspondence between ***c*** and ***t*** such that *c*_1_ = *t*_1_, *c*_2_ = *t*_2_ etc. For *G*_2_ and *G*_3_, ***c*** is a permutation of the times ***t***.

| Case | $(\tau_1, \tau_2)$ | $(\tau_2, \tau_3)$ | $(\tau_3, \tau_4)$ | $> \tau_4$ |
| --- | --- | --- | --- | --- |
| 1 | $c_1$ | $c_2$ | $c_3$ | |
| 2 | $c_1$ | $c_2$ | | $c_3$ |
| 3 | $c_1$ | | $c_2, c_3$ | |
| 4 | $c_1$ | | $c_2$ | $c_3$ |
| 5 | $c_1$ | | | $c_2, c_3$ |
| 6 | | $c_1, c_2$ | $c_3$ | |
| 7 | | $c_1, c_2$ | | $c_3$ |
| 8 | | $c_1$ | $c_2, c_3$ | |
| 9 | | $c_1$ | $c_2$ | $c_3$ |
| 10 | | $c_1$ | | $c_2, c_3$ |
| 11 | | | $c_1, c_2, c_3$ | |
| 12 | | | $c_1, c_2$ | $c_3$ |
| 13 | | | $c_1$ | $c_2, c_3$ |
| 14 | | | | $c_1, c_2, c_3$ |

### 3.2 Site pattern probabilities given gene trees

The calculation for site pattern probabilities given the gene tree is the standard probability calculation one would perform absent any consideration of the coalescent process [5]. From the Jukes-Cantor model, we have that when time is measured in mutation units the transition probabilities on branch *l* with length *d*_*l*_ are given by:

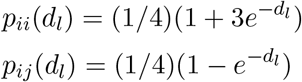

where *i, j* ∈ {*A, C, G, T* }.

The probability of the states of the gene tree displayed in figure 3 is the probability of any state at one of the interior nodes (which is 1/4 for all four nucleotides) times the transition probability over the seven branches. It is for this reason we treat the gene trees as unrooted: it gives us one fewer branch substitution probability to concern ourselves with. The nucleotide at the root can be safely ignored because the Jukes-Cantor model is a subset of the general time reversible (GTR) model, and so by the Law of Total Probability and the time-reversibility of this time-homogeneous stochastic process the probability of any state of the tree (excluding the root) is the probability of the state given the nucleotide at the root times the probability of that nucleotide (1/4 under the JC69 assumptions) summed over all four possible nucleotides. Then, the probability of one particular state of the tree (including knowledge of the state of the interior nodes) is given by:

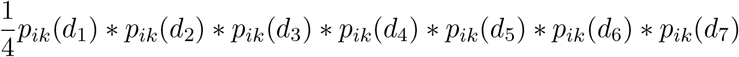

where *k* ∈ {*i, j*}. To find the probability of a given site pattern, we perform an exhaustive search over all node states, identify which states fit which site pattern, and sum common terms. The results are implemented in our C++ code available on GitHub.

**Figure 3:**
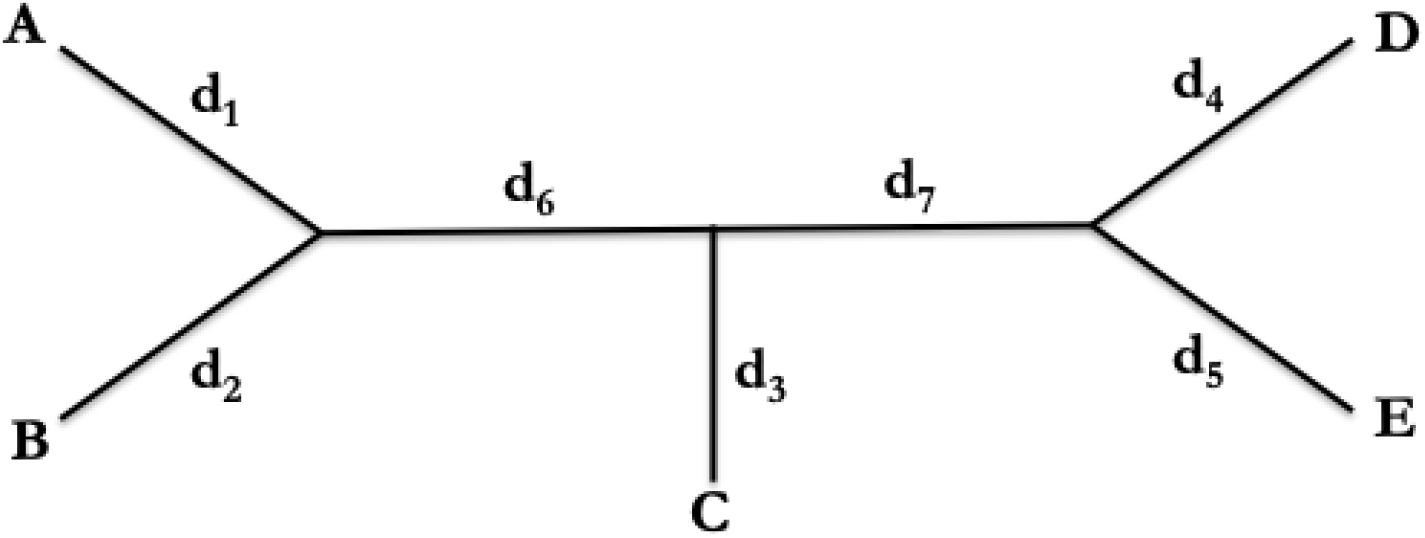
Example unrooted 5-taxon tree with distances

We now note that each of the terms *p*_*ik*_(*d*_1_) ∗ *p*_*ik*_(*d*_2_) ∗ *p*_*ik*_(*d*_3_) ∗ *p*_*ik*_(*d*_4_) ∗ *p*_*ik*_(*d*_5_) ∗ *p*_*ik*_(*d*_6_) ∗ *p*_*ik*_(*d*_7_) is a linear combination of 128 terms of the form:

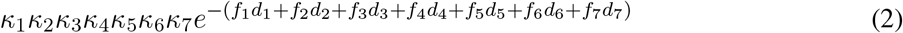

where each of the *κ*_*m*_ ∈ {−1, 3} and *f*_*m*_ ∈ {0, 1} (and each term is multiplied by 4^−7^).

Taken together, each of the 51 different site pattern probabilities given the gene tree is a linear combination of the terms in equation 2. The result is another linear combination (the coefficients are calculated by our C++ code):

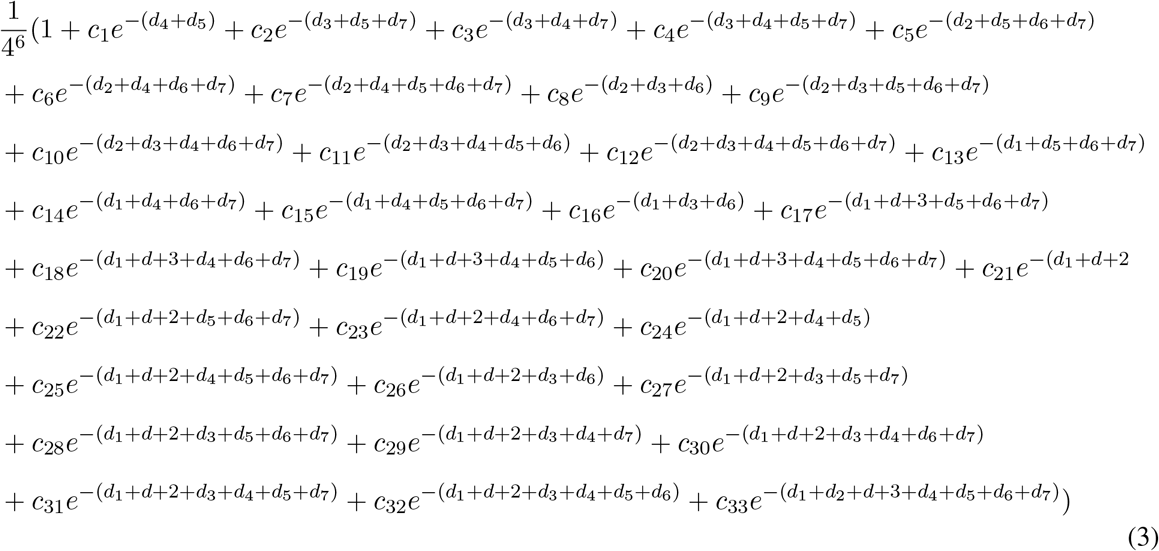

Next, we need to express the terms in equation 3 in terms of our rates, speciation times and coalescent times. Consider the example shown in figure 2. Putting the distances on the tree in coalescent units, assuming that *α* = 4*/*3 as specified by the Jukes-Cantor model, and adding in the factor *θ/*2 to account for the conversion from mutation to coalescent units, we have that:

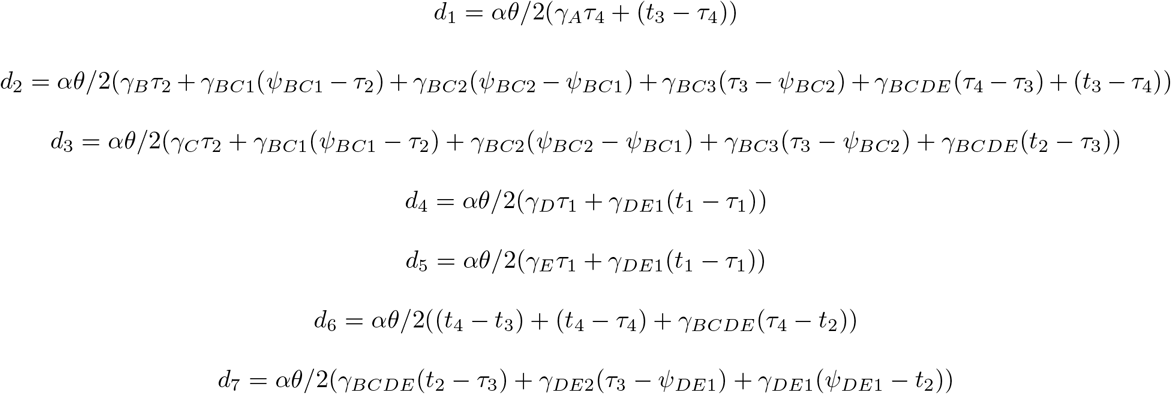

Then, we note that each term of equation 3 can be written in the form 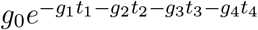. For example, for the example shown in figure 2, and rearranging terms

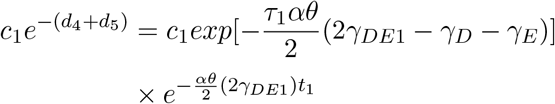

So 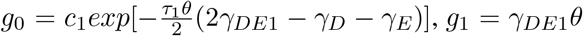, and *g*_2_ = *g*_3_ = *g*_4_ = 0. Similarly, we can rewrite all the terms of equation 3 where *g*_0_ is the sum of *αθ/*2 multiplied by the time of each of the break points that occur along the branch multiplied by the difference in the relative mutation rates before and after the break point, or *ψ*_*j,i*_(*γ*_*j,i*−1_ − *γ*_*j,i*_). (In the event that *i* = 0 we can define *γ*_*j,i*−1_ to be the final rate of the branch below *j*.) We can further show that the *g*_1_, *g*_2_, … constants are equal to 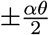 times the rate at that coalescent point if the branch is not included in the term of equation 3, with the sign being positive if the branch descends from the coalescent point to a leaf and negative if the branch ascends from the coalescent point to the root. The constant is zero if the branch is not included in that term of equation 3.

### 3.3 Gene tree probabilities given the species tree

#### 3.3.1 Background

The gene tree density for a given gene tree arising within a species tree is given by [17] and [2]. Given *j* lineages, and time measured in coalescent units, the time to the next coalescent event *t*_*j*_ follows an exponential distribution with rate 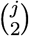. For any interior branch of the species tree, this process is censored, and so with *j* lineages entering a branch of length *τ*_*b*_ the probability of no coalescent events on the branch is given by 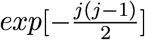. More generally, the density of the vector of coalescent times for which *j* lineages enter an interior branch and *k* leave is given by:

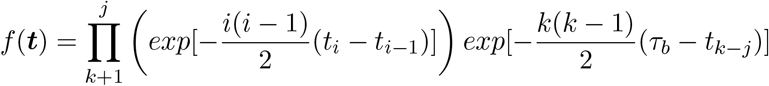

where *t*_0_ = 0.

For example, given the species and gene trees in figure 2, we see *B* and *C* do not coalesce on their first common branch, which occurs with probability 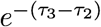. *D* and *E* coalesce on their first common branch with a density 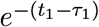. Three lineages enter the common *BCDE* branch and two leave, which has a density 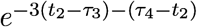. Three lineages enter above the root, so the density of the final coalescent events is 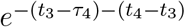. Taken together, the probability of this gene tree is:

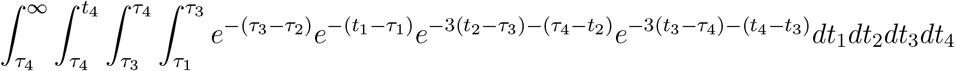

Then, combining with the results from section 3.2 and rearranging terms we have that the probability of the data given this gene tree history and species tree is a linear combination terms of the form:

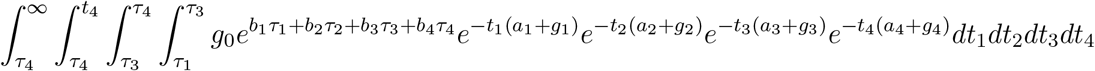

where in this case ***b*** = (1, 1, 2, 2) and ***a*** = (1, 2, 2, 1).

More generally, the different integrals that need to be evaluated depend on several factors. First, we need to consider the limits of integration. Specifically, the form of the integrals will change depending on whether multiple coalescent events occur within the same branch and if they are separated by a rate change. We introduce the notation 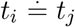 to indicate that these two coalescent events occur in the same branch and rate category and 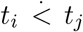 to indicate they occur in different branches and/or have a rate change between them. We need to consider eight cases:

- 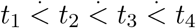
- 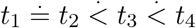
- 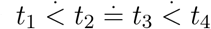
- 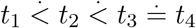
- 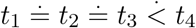
- 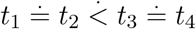
- 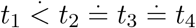
- 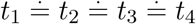

For each of the cases in table 2, the limits of integration will change, as will the vectors ***b*** and ***a***. Further, for cases where multiple coalescent events occur on the same branch, we must be able to distinguish whether or not the coalescent event occur on the same side of any rate change breakpoints.

Our implementation first evaluates each of the eight enumerated integral forms in general terms as a function of the eight limits of integration, and the vectors ***g, b*** and ***a***. Next, it searches over all possible gene tree histories, determines the applicable integral form, finds the ***g*** as in section 3.2, and applies the appropriate ***b*** and ***a*** based upon the gene tree history. These probabilities are then summed to find the final site pattern probability given the species tree.

## 4 Implementation

Four programs, all written in C++ are provided at https://github.com/richards-1227/fiveTaxa. A single C++ file is included as well as a Mac and Unix executable. Preliminary work, such as the enumeration of gene tree histories, calculation of the *c*_*i*_ terms in equation 3, and defining the permutation matrix are commented out at the end of the C++ file, but the interested reader can use the code to follow our steps as interested.

Execution is via the command

~~~
./Taxa5 topInd fileName *θ*
~~~

topInd = 0, 1, or 2 for *S*_1_, *S*_2_, and *S*_3_, respectively.

“fileName” is the name of a text file containing the sets of rates and breakpoints in the following format:

~~~
branch start_time end_time rate
~~~

For example, the tree in figure 2 would have topInd = 1 and the following input in fileName:

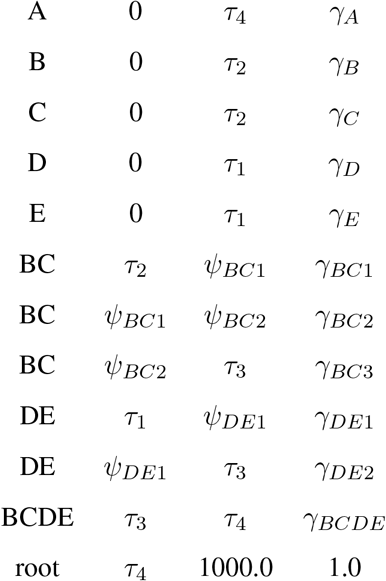

The final time above the root must be chosen such that *e*^*ψ*^ ≈ 0. We have used 1000.0.

To check the accuracy of the implementation, we first generated site pattern probabilities for different branch lengths with ***γ*** equal to a vector of ones, representing a strict molecular clock. We compared that output to existing code which implements the results of [2] and confirmed that the results were identical after marginalizing over each taxon in turn. We then implemented up to four different rates simultaneously on each branch of the tree, varying the rates by up to a factor of 16, and compared the results to those derived by [19], again marginalizing over each taxon in turn and verified the results matched.

## 5 Applications

Here, we introduce SVDquintets, a new method for species tree inference under the multispecies coalescent model. Chifman and Kubatko (2015) introduced the SVDQuartets algorithm, and much of this discussion follows from that paper, with adjustments as needed for five taxa. We define a *split* as any bipartition of the leaf set *L* into *L*_1_ and *L*_2_. We say the split is *valid* if the removal of an edge in *S* can produce two subtrees *S*_1_ and *S*_2_ such that the leaf set 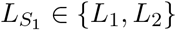. Let a *cherry* refer to a pair of species more related to each other than to any other species in *L*. Then *L*_2_ being a cherry represents a special case of a valid split where |*L*_2_| = 2.

We next define a flattening matrix **D**_*flat*_. This matrix is a 4^3^ ×4^2^ rearrangement of site pattern frequencies where the rows represent the possible states {*A, C, G, T* }^3^ = {*AAA, AAC, AAG*, … *TTG, TTT* } of the taxa in *L*_1_ and the columns represent the possible states {*A, C, G, T* }^2^ of the taxa in *L*_2_. So, for example, the the entry corresponding to the *ACC* row and the *GT* column is the number of sites with the pattern *ACCGT*. Finally, let ***δ***_*flat*_ refer to the set of probabilities associated with each element of ***D***_*flat*_ – so if *L*_1_ = {*A, B, C*} and *L*_2_ = {*D, E*}, *δ*_*AAA*|*T C*_ = *P* (*i*_*A*_ = *A, i*_*B*_ = *A, i*_*C*_ = *A, i*_*D*_ = *T, i*_*E*_ = *C*) at any given site. With five taxa, we can form ten different flattening matrices for the 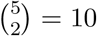 possible subsets of taxa in *L*_2_.

Just as we can form a flattening matrix of the observed data **D**_*flat*_, we can form a flattening matrix of the site pattern probabilities ***δ***_*flat*_, ordered in the same manner. Chifman and Kubatko (2015) demonstrated that for four taxa the rank of the flattening matrix *rank*(***δ***_*flat*_) ≤ 10 if and only if the split is valid. For the observed data, we can perform a singular value decomposition ***D***_*flat*_ = ***U* σ*V*** ^*T*^ where ***U*** and ***V*** are orthogonal matrices and **σ** is a diagonal matrix of singular values (ordered from greatest to least in magnitude). Then we can then take the sum of the highest order singular values as a measure of how “close” the split is to having rank ≤ 10. For a given split, we define its SVD score as 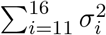 where *σ*_*i*_ are the ordered singular values, and by the law of large numbers the SVD score will approach zero asymptotically as the number of sites goes to infinity. We will demonstrate, at least empirically, that this result also appears to hold for five taxa under the assumptions of our data generating model.

Finally, we define a second split *L*_3_|*L*_4_ as *compatible* with the first split *L*_1_|*L*_2_ if *L*_4_ ⊂ *L*_1_. For example, let *L*_1_ = {*A, B, C*} and *L*_2_ = {*D, E*}. Then *C, D, E*|*A, B* would be a compatible split but *A, C, E*|*B, D* would not be. Note that for five taxa there are three second splits compatible with any given first split, and that any set of two compatible splits defines an unrooted tree. We then propose the following algorithm for inferring an unrooted 5-taxon species tree:

### Algorithm 1

SVDquintets

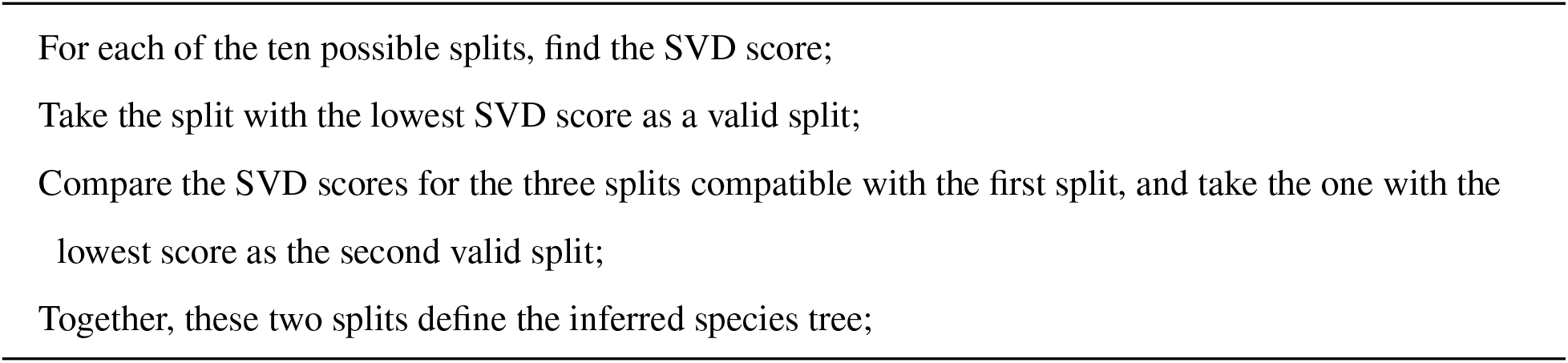

Because we do not have exact algebraic expressions for the site pattern probabilities, we simulated various species trees, and then calculated the site pattern probabilities given these species trees. We then formed flattening matrices given these calculated probabilities, and compared the SVD scores of the various flattening matrices for valid and invalid splits.

The simulation settings are as follows: speciation times were sampled following the Yule model, with *τ*_1_ ∼ *Exp*(3*/*5), (*τ*_2_ − *τ*_1_) ∼ *Exp*(3*/*4), (*τ*_3_ − *τ*_2_) ∼ *Exp*(3*/*3), and (*τ*_4_ − *τ*_3_) ∼ *Exp*(3*/*2), where here the exponential parameter refers to the mean waiting time. These settings were chosen so that on average the root would be approximately 4 coalescent units from the present. The mutation rate above the root was set at unity, and mutation rates on pendant branches were set at 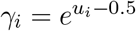 where 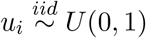. Mutation rates on interior branches were set as the arithmetic mean of the pendent branches descending from them. *θ* was set following a uniform distribution (on the log scale) between 0.0003 and 0.03.

One thousand trees were simulated for each of *S*_2_ and *S*_3_. For *S*_1_, 400 trees were simulated for each of the cases *τ*_3_ *< τ*_1_, *τ*_1_ *< τ*_3_ *< τ*_2_ and *τ*_2_ *< τ*_3_ for a total of 3200 simulations. Because the site pattern probabilities are the sum of a large number of very small values, we performed all our calculations with long double (128-bit) precision. Nevertheless, when the rank of the flattening matrices were calculated, none of the SVD scores were exactly zero, which we suspect is due to very small floating point errors arising in the calculation of probabilities and the subsequent matrix decomposition. There was, however, a noticeable difference in SVD scores for valid and invalid splits, suggesting that the algorithm is useful for inference. Further, in all 3200 cases, the minimum SVD score was from a valid split, and the minimum SVD score of a compatible second split was the valid second split. The data are summarized in table 3:

**Table 3:** Results for SVD scores for 3200 simulated trees. The second column gives the average minimum SVD score for a valid split. The third column gives the average ratio between the lowest SVD score of a valid and invalid split. The last column gives the ratio between the SVD score of the valid second split to the lowest SVD score of an invalid split compatible with the first split.

| Topology | Avg min valid score | Avg ratio valid/invalid | Avg ratio valid/invalid – second |
| --- | --- | --- | --- |
| $S_1$ | $3 \times 10^{-8}$ | $1.4 \times 10^8$ | $1.36 \times 10^7$ |
| $S_2$ | $3 \times 10^{-8}$ | $1.4 \times 10^{10}$ | $4.7 \times 10^9$ |
| $S_3$ | $3 \times 10^{-8}$ | 300 | 400 |

We next tested the SVDquintets algorithm against SVDQuartets using simulated data. 300 trees were simulated for each topology as above, except for each setting 100 trees were simulated with an expected root age of 4 coalescent units, 100 with an expected root age of 2 coalescent units, and 100 with an expected root age of 1 coalescent unit. 100,000 sites were simulated for each tree. The mean RF distance [20] between the estimated and true tree was calculated, as well as the mean RF distance when both SVDquintets and SVDQuartets estimated the same tree.

We can see from table 4 that SVDquintets shows improvement over SVDQuartets with topologies *S*_1_ and *S*_2_, and does no worse with *S*_3_. (The slight difference between the two in *S*_3_ arises from the SVDQuartets implementation in PAUP* [25] which returns a non-binary tree for some difficult cases, whereas our SVDquintets implementation will always return a binary tree.) When both methods agree, the estimated tree is even more accurate on average than any individual method, although the frequency with which the two methods agree can be low for difficult settings, especially with *S*_2_ or *S*_3_.

**Table 4:** Mean RF distance from the estimated to the true tree for SVDQuartets and SVDquintets. The number after the dash indicates the expected root age. The ensemble row refers to cases when the two methods agreed on the inferred tree. The percentage of times this occurred is given in parentheses.

| Method | $S_1 - 4$ | $S_1 - 2$ | $S_1 - 1$ | $S_2 - 4$ | $S_2 - 2$ | $S_2 - 1$ | $S_3 - 4$ | $S_3 - 2$ | $S_3 - 1$ |
| --- | --- | --- | --- | --- | --- | --- | --- | --- | --- |
| SVDQuartets | 0.29 | 0.41 | 0.79 | 0.22 | 0.84 | 1.31 | 0.46 | 0.93 | 1.49 |
| SVDquintets | 0.22 | 0.34 | 0.64 | 0.16 | 0.50 | 1.08 | 0.46 | 0.98 | 1.50 |
| Ensemble | 0.15(91) | 0.24(90) | 0.59(82) | 0.02(83) | 0.00(54) | 0.51(39) | 0.22(80) | 0.69(75) | 0.96(48) |

## 6 Conclusions and future work

This work has a number of advantages. First, we can directly compute likelihoods for five-taxon trees under a variety of time-varying rate assumptions. Second, we have introduced a new species tree estimation method that appears to work better than its closest existing analog under various simulated conditions.

This work may be able to be generalized further. First, this general format for finding site pattern probabilities can be extended to six or more taxa, but may be limited by computational challenges.

More work also needs to be done to verify that the advantage seen here for SVDquintets continues to hold. We would expect a method that treats larger numbers of taxa simultaneously would be more accurate as it can account for the covariances in site pattern probabilities in a way a four-taxon method cannot. But, conversely, as you increase the number of taxa, the probability of the least probable site patterns decreases (the probability of XYZWX must necessarily be less than the probability of XYZW). As a result, ever more data is required with more taxa to prevent sparsity of the flattening matrix to erode or reverse whatever advantage is gained from jointly estimating the topology of higher numbers of taxa. Additionally, there is already an extensive literature on inferring larger trees using quartet inputs; we would have to develop similar inferential methods using quintet inputs.

Extending beyond five taxa faces the problem that not all splits are the same size. With five taxa, we have two 3,2 splits in the unrooted tree. But with six taxa you either have three 4,2 splits or two 4,2 splits and a 3,3 split. Allman et al. (2017) demonstrated that splits of different sizes may not be comparable using SVD scores, so another approach may need to be taken to perform estimation of higher taxa trees using ranks.

## 7 Acknowledgments

The author wishes to thank Dr. Laura Kubatko for her suggestions and support through this work.

